# Clade B serpins promote direct basal-to-goblet cell differentiation in airways of patients with chronic obstructive pulmonary disease

**DOI:** 10.64898/2026.09.21.753278

**Authors:** Tiffany S. Tufenkjian, Jessica B. Blackburn, David S. Nichols, Alfredo Vasquez, Ciara M. Shaver, Lorraine B. Ware, Timothy S. Blackwell, Bradley W. Richmond

## Abstract

Goblet cell hyperplasia (GCH) is a hallmark of chronic obstructive pulmonary disease (COPD) and contributes to morbidity and mortality. We investigated the cellular and molecular origins of GCH in COPD using single-cell RNA sequencing (scRNA-seq), spatial transcriptomics, and in vitro models. We identified “basal-to-goblet transitional cells” (BGTC) which are transcriptionally and physically located between basal and goblet cells and are characterized by expression of clade B serpins. In vitro studies indicate that inflammatory cytokines, including IL-1β, stimulate basal-to-goblet differentiation through a *SERPINB3/4*^+^ intermediate and *SERPINB3* overexpression induces differentiation of airway basal cells into *IL1B*-producing inflammatory goblet cells. In addition, we identified a subset of *SERPINB4*^+^ “goblet-variant basal cells” from lungs of COPD patients that express goblet cell and inflammatory genes ex vivo. Together, these results indicate clade B serpins contribute to altered epithelial differentiation and inflammatory signaling in COPD and support therapeutic targeting of clade B serpins to reduce GCH.

## INTRODUCTION

The airway epithelium is comprised of multiple specialized cell types that function in a coordinated manner to protect against inhaled microbes and non-infectious irritants (1, 2). Within a species, proportions of these cells are relatively fixed in similarly-sized airways. However, in the setting of acute infection or exposure to inhaled irritants, mucus-producing goblet cells expand in number, facilitating entrapment of microbes or inhaled irritants at the cost of regional defects in gas exchange due to impaired mucociliary clearance and mucus plugging. Expansion in goblet cells involves activation of SAM pointed domain-containing Ets transcription factor (SPDEF) by inflammatory cytokines including IL-4, IL-13, IL-17, and IL-Iβ (3–6), thereby coupling inflammation to goblet cell hyperplasia (GCH). This coupling ensures goblet cells return to their homeostatic proportions once the inciting infection or irritant is removed, and inflammatory cytokines return to baseline levels.

Chronic obstructive pulmonary disease (COPD) is a heterogeneous lung condition characterized by persistent expiratory airflow obstruction and caused by impaired lung growth, accelerated lung function decline, or both (7, 8). Chronic exposure to cigarette smoke or other inhaled irritants at all life stages is a major risk factor (9–11). While current therapies improve symptoms and reduce hospitalizations, COPD remains the third leading cause of death from chronic disease worldwide (12). GCH is a hallmark of COPD and contributes to airflow obstruction directly through formation of viscous mucus plugs (13) which are associated with disease exacerbations, impaired lung function, and increased mortality (14–16). Additionally, we have shown that GCH is associated with a local reduction of secretory immunoglobulin A (SIgA) which contributes to ongoing small airway inflammation and fibrosis (14, 15, 17–19).

Despite the importance of GCH to COPD, the origin of goblet cells in the COPD lung remains unclear. In vitro and in vivo studies indicate goblet cells may arise via trans-differentiation from other airway epithelial cell sub-types, particularly secretory cells, in response to COPD-relevant stimuli such as cigarette smoke, viral or bacterial infection, neutrophil elastase, and hypoxia (20–26). In contrast, other studies have indicated that some COPD basal cells are primed toward goblet cell differentiation, suggesting direct differentiation from airway stem cell progenitors (22, 27, 28). While a prior single-cell RNA sequencing (scRNA-seq) study of the human COPD lung suggested goblet cells arise primarily from secretory (club) cells (29), a paucity of basal cells in this study limited a full assessment of potential basal-to-goblet cell differentiation. Given the critical role of GCH in COPD, a better understanding of how goblet cells form in the COPD lung is needed to develop new therapeutic targets.

Here, we coupled scRNA-seq and spatial transcriptomic analyses of human COPD lung tissue with in vitro models to determine the cell of origin of goblet cells in COPD and identify novel disease-related pathways for therapeutic targeting. Our results indicate goblet cells form from basal cells in COPD via an intermediate cell state that responds to and produces inflammatory cytokines via clade B serpins.

## RESULTS

### Basal-to-goblet transitional cells (BGTCs) are enriched in areas of GCH in COPD lungs and express clade B serpins

To investigate how goblet cells form in human lungs with COPD, we used our previously curated single-cell RNA sequencing (scRNA-seq) atlas (30) that included distal lung samples from COPD patients who underwent lung transplantation at multiple centers (n=22) and deceased donors without COPD whose lungs were rejected for transplantation (i.e. controls, n=18). We focused our attention on the non-ciliated airway epithelial cell types (30), which included basal, goblet, and secretory cells (including *SCGB1A1*^+^ secretory and *SCGB1A1*^+^*SCGB3A2*^+^ respiratory airway secretory cells or RASCs) and a cluster of squamous cells unique to COPD samples (**Fig. 1A**). Goblet cells were abundant in samples from COPD patients and located near basal cells on a Uniform Manifold Approximation and Projection (UMAP) plot (**Fig. 1A**). Additionally, we observed rare cells co-expressing the basal cell marker KRT5 and the goblet cell marker MUC5B in COPD airways by immunostaining (**Fig. S1A**), suggesting a possible basal-to-goblet intermediate cell. To investigate this further, we sub-clustered basal and goblet cells. This analysis revealed a population of cells originally annotated as goblet cells that was almost entirely COPD-specific and localized to a transitional area between basal cells and goblet cells on the original UMAP (**Fig. 1B, Fig. S1B**). We annotated these cells “basal-to-goblet transitional cells” or BGTCs. BGTCs expressed the basal cell genes *TP63, KRT5, ITGB4* and the goblet cell transcription factors *SPDEF* and *XBP1* but did not express the mucins *MUC5AC* or *MUC5B* (**Fig. 1C**). To further characterize gene expression in these cells, we performed a gene set enrichment analysis based on gene ontogeny (GO) in BGTC marker genes. Compared to other airway epithelial cell types, including multiciliated cells (MCCs), BGTCs had reduced expression of genes related to ciliogenesis and increased expression of genes related to carbohydrate, organic acid, and nucleotide metabolism, energy production, and cell adhesion and migration (**Fig. 1D**). Interestingly, a top activated pathway (normalized enrichment score of 2.72, adjusted p = 8.51 x 10^-6^) was negative regulation of endopeptidase activity (GO:0010951) which includes the clade B serpin family (**Table S1**). Multiple members of this family including *SERPINB1, -B2, -B3, -B4, -B5, -B6, -B7, -B8, -B10, - B11,* and *-B13* were expressed by BGTCs (**Fig. S1C**).

**Fig. 1.**
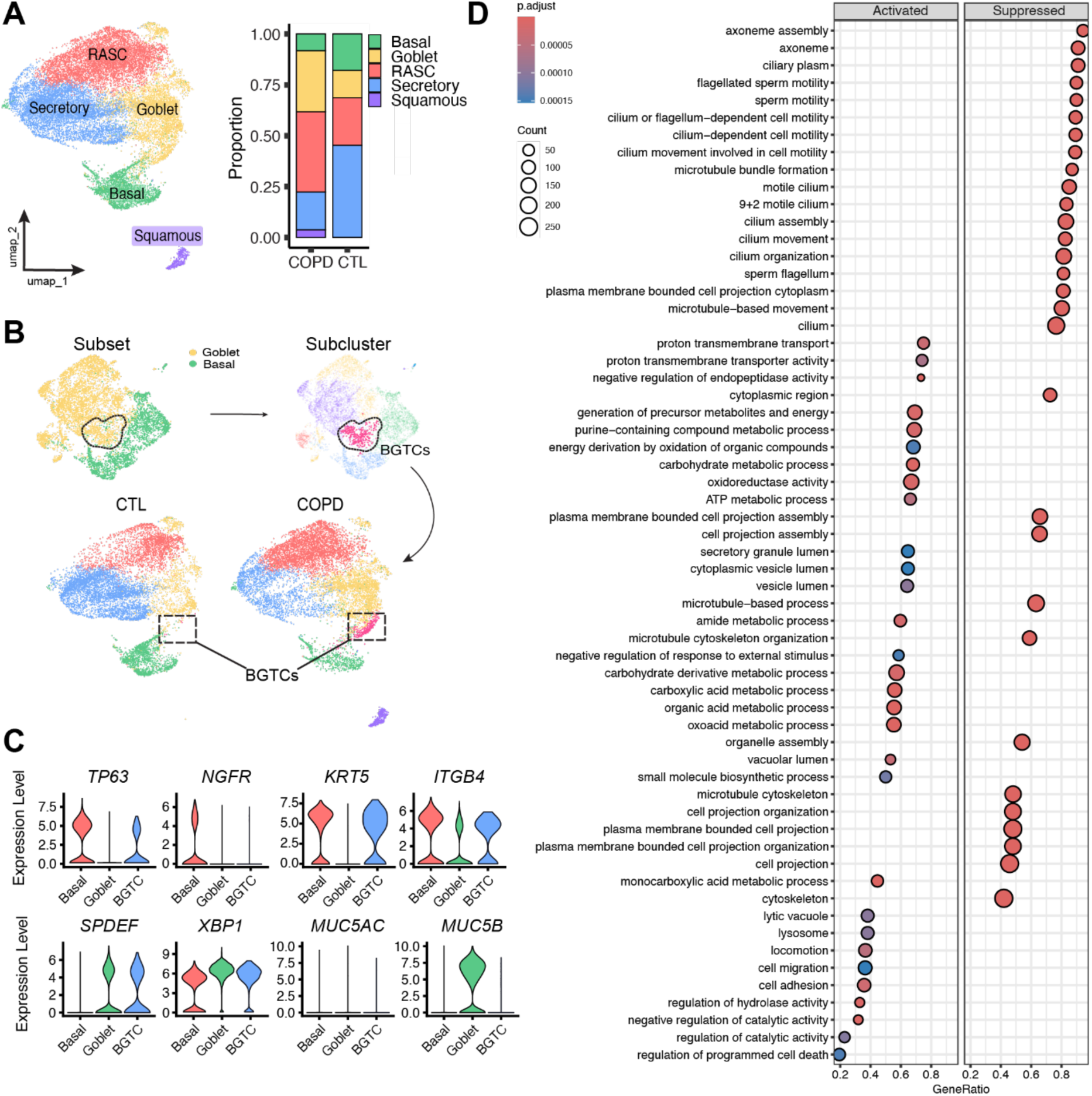
Basal-to-goblet transitional cells (BGTCs) are enriched in COPD lungs and express clade B serpins. **(A)** Uniform manifold projection (UMAP) of the small airway epithelial cell (SAEC) object, excluding ciliated cells, from COPD and control donors. Proportion plot of each cell type in COPD and control. **(B)** Sub clustering of basal and goblet cells showing COPD-enriched BGTC population, and its location on the original UMAP **(C)** Expression of basal cell markers (*TP63, NGFR, KRT5* and *ITGB4*) and goblet cell markers (*SPDEF, XBP1, MUC5AC,* and *MUC5B*) in Basal, Goblet, and BGTC annotated cell types. **(D)** Gene set enrichment was performed on BGTC markers for GO biological process terms. suppressed pathways are those that are down in BGTCs while activated pathways are up. The size of the dot represents the number of genes in that term while the color represents adjusted p values.

To localize BGTCs in intact human lung tissue, we first utilized an RNA in situ hybridization probe targeting the clade B serpins *SERPINB3* and *SERPINB4,* which share 98% nucleotide and 92% amino acid sequence homology (31). We observed substantial expression of *SERPINB3/4* in COPD airways with MUC5B^+^ goblet cells, but little expression in control airways without GCH (**Fig. 2A**). To investigate this further, we developed a custom probe panel for the Xenium spatial transcriptomics platform (10X Genomics) targeting the top 50 markers of BGTCs derived from our scRNA-seq atlas including *SERPINB3*, -*B4*, -*B5*, and -*B13* and other genes of interest (**Fig. 2B, Table S2**). This panel was used alongside the Xenium human lung panel for cell type annotation, and applied to formalin-fixed, paraffin-embedded (FFPE) lung sections from two COPD lung explants and two deceased donor controls to annotate all major cell types (**Fig. S2A)**. Unsupervised clustering of non-alveolar epithelial cells (**Fig. S2B**) showed BGTCs were enriched in COPD lungs and located between basal and goblet cells on a UMAP plot (**Fig. 2C**) similar to what we observed in our scRNA-seq atlas. We mapped these clusters back onto H&E-stained sections and found that BGTCs were physically located between basal and goblet cells and largely restricted to COPD airways with GCH (**Fig. 2D**). *SERPINB3/4* was expressed at low levels in basal cells from COPD and control airways but was much higher in BGTCs and some goblet cells (**Fig. 2D**), consistent with our observations using RNA in situ hybridization (**Fig. 2A**).

**Fig. 2.**
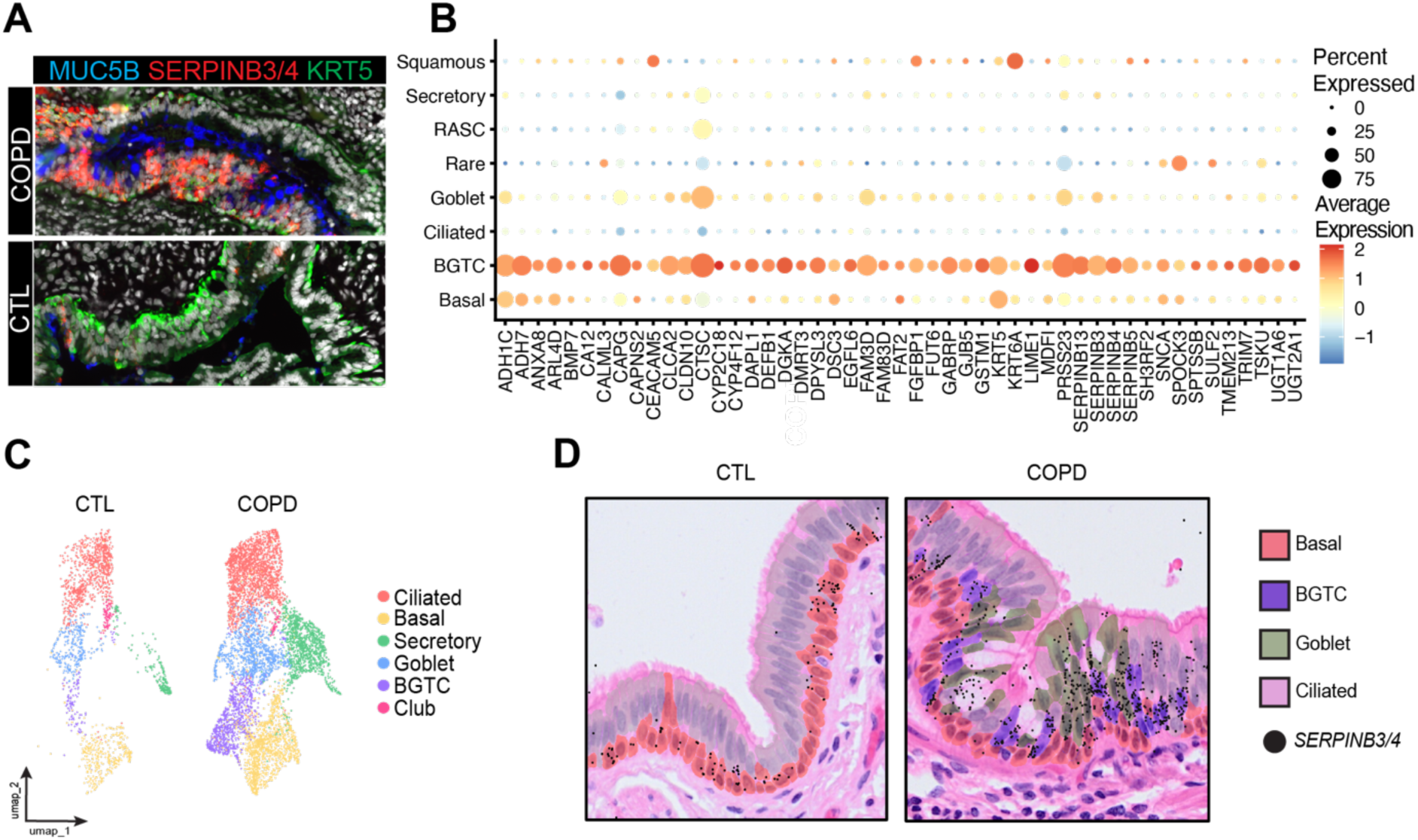
Basal-to-goblet transitional cells (BGTCs) are enriched in COPD lungs in regions of GCH and express clade B serpins. **(A)** Immunostaining and RNA-ISH in control and COPD airways. MUC5B (blue), *SERPINB3/4* (red), and KRT5 (green). **(B)** Dot plot of top 50 markers in BGTC cells compared to all other SAECs. **(C)** UMAP of airway cell types from control and COPD lungs profiled with Xenium spatial transcriptomics. **(D)** Xenium airway annotations and localization/expression of *SERPINB3/4* transcripts overlaid on post-Xenium H&E staining of control (left) and COPD (right) airways.

Together, these results indicate a basal-to-goblet cell transitional state characterized by high expression of clade B serpins in addition to genes related to metabolism, energy production, and cell adhesion and migration. These cells are highly enriched in regions of GCH in the lungs of patients with COPD. Although rarely observed in non-diseased lungs, BGTCs when present, are accompanied by goblet cells.

### Basal cells differentiate into goblet cells in response to cytokine stimulation through an intermediate state characterized by clade B serpin expression

Next, we investigated how goblet cells form in vitro in response to cytokine stimulation. We chose to induce GCH with IL-1β and IL-13 given their known ability to generate goblet cells in vitro and their importance in COPD and asthma, respectively (26). Primary human small airway epithelial cells (HSAECs) from non-COPD controls (n=3) were treated with IL-1β or IL-13 during days 14-27 of air-liquid interface (ALI) culture. IL-13-induced goblet cells had a classic goblet cell morphology when viewed cross-sectionally on Alcian Blue Periodic acid-Schiff (AB-PAS)-stained membrane sections and expressed high levels of MUC5AC, while IL-1β-induced goblet cells were dysmorphic and expressed high levels of MUC5B (**Fig. 3A**). We used split-pool ligation-based sequencing (SPLiT-seq) (32) to perform scRNA-seq on ALI-differentiated cells from each group. After recursive clustering we annotated nine non-ciliated airway epithelial cell types based on canonical markers including resting and proliferating basal cells; squamous basal cells; intermediate and mature secretory cells; and intermediate and mature IL-1β and IL-13-induced goblet cells (**Fig. 3B, Fig. S3A and B, Table S3**).

**Fig. 3.**
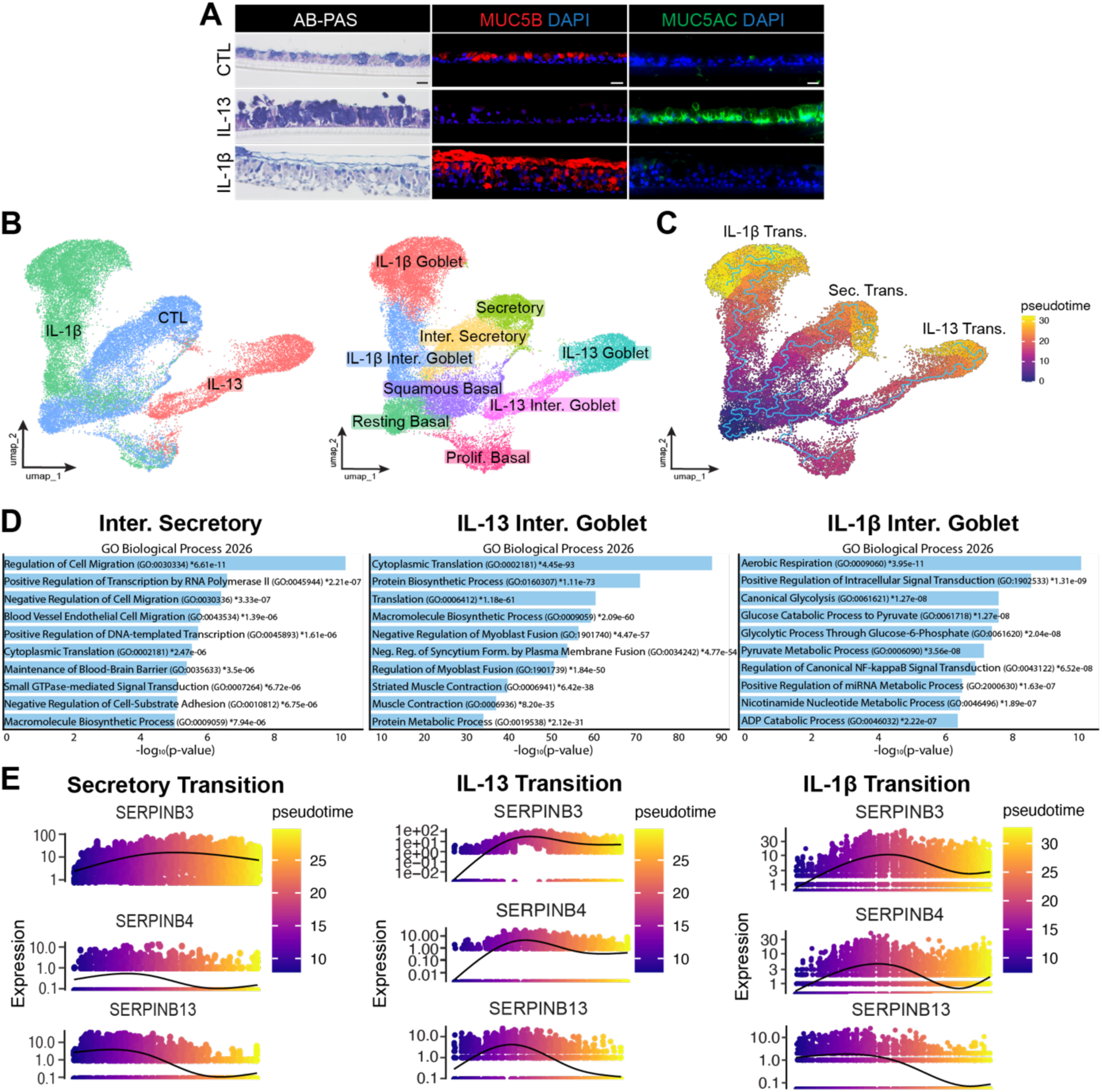
Basal cells differentiate into goblet cells via an intermediate population characterized by clade B serpin expression in vitro in response to IL-13 or IL-1β. Human small airway epithelial cells (HSAECs) were differentiated in air-liquid interface (ALI) culture for 29 days (D29) and exposed to IL-1β (10 ng/mL) or IL-13 (10 ng/mL). **(A)** Representative images of ALI membranes in cross-section for each treatment group collected on ALI D28. Alcian Blue-Periodic Acid-Schiff (AB-PAS) staining of mucin positive cells (left panel). Immunostaining for Mucin 5b (MUC5B, red, middle panel) and Mucin 5ac (MUC5AC, green, right panel) with DAPI stained nuclei (blue). Scale bar = 20 μM. **(B)** UMAP of cytokine treated cells or controls colored by treatment group (left) UMAP colored by annotated cell type (right). **(C)** Pseudotime plotted on the original UMAP using Monocle3 with resting basal cells set as time zero. **(D)** Enrichr bar plots based on cell type markers showing top GO Biological Process terms (2026) for each intermediate secretory cell population in **C**. **(E)** Gene expression over pseudotime plots of SERPINB3, SERPINB4, and SERPINB13 for each of the three secretory branches shown in **D**.

IL-1β and IL-13-treated cells formed distinct clusters of cells (**Fig. 3B**) marked by high expression of *MUC5B* and *MUC5AC*, respectively (**Fig. S3B**), similar to what was observed by immunostaining (**Fig. 3A**). A trajectory analysis performed using Monocle 3 (33, 34) indicated both IL-1β and IL-13-treated goblet cells primarily arose from basal cells rather than mature secretory cells (**Fig. 3C**). While cells transitioning between basal and mature secretory cells in non-inflamed conditions showed enrichment for cell migration and positive regulation of transcription when analyzed by GO using Enrichr (35, 36) (**Fig. 3D, left panels**), we found different transcriptional patterns in intermediate cells transitioning toward goblet cells in response to IL-13 or IL-1β . While cells transitioning between basal and goblet cells in response to IL-13 (IL-13 intermediate goblet cells) showed enrichment in terms related to translation and protein synthesis (**Fig. 3D, middle panels**), cells transitioning between basal and goblet cells (IL-1β intermediate goblet cells) showed enrichment in aerobic respiration and glycolysis (**Fig. 3D, right panels)**. Interestingly, many of the top terms in IL-1β-treated cells were similar to those enriched in BGTCs (**Fig. S3C**).

Since increased expression of clade B serpins was observed in BGTCs (**Fig. S1C**), we next examined expression of *SERPINB3*, -*B4*, and -*B13* over pseudotime in IL-1β and IL-13-treated cells and untreated cells. Generally, we noted increased expression of these serpins as basal cells differentiated into goblet cells in response to cytokine stimulation and in untreated basal cells as they differentiated into mature secretory cells (**Fig. 3E**). Together, these data show goblet cells differentiate primarily from basal cells in response to inflammatory cytokines and that clade B serpins are transiently increased during basal-to-goblet cell differentiation in vitro. **SERPINB3 overexpression induces GCH in vitro**

We next sought to externally validate whether clade B serpins and goblet cell genes are co-expressed in human lungs. To accomplish this, we examined gene expression correlations between *SERPINB3*, *B4*, or *B13* in human scRNA-seq from the Human Lung Cell Atlas (HLCA) using LungChat (37). In resting basal cells, *SERPINB3*, -*B4*, and -*B13* were highly correlated (Pearson r = 0.795 for *SERPINB3* and *B13*; r = 0.778 for *SERPINB3* and *B4*, and r = 0.696 for *SERPINB4* and *B13*). Using Azimuth-based cell annotations through Enrichr, (36) we queried the cell type most closely associated with *SERPINB3*, -*B4*, or -*B13*-associated genes. Goblet cells were the top associated cell type for *SERPINB3* and -*B4* while proximal basal cells were the top associated cell type for *SERPINB13* (goblet cells were second) (**Table S4, Fig. S4A**).

Interestingly, a shared set of 67 genes associated with *SERPINB3*, -*B4*, and -*B13* included multiple gene associations with squamous cell genes from our scRNA-seq atlas (**Fig. 1A**) including *IGFBP3*, *S100P*, *PMAIP1*, *KRT6A*, *CLDN1*, *TPM4*, and *EMP1* (**Fig. S4B)** suggesting clade B serpins might also be associated with squamous cell differentiation.

To directly test whether clade B expression in basal cells alters epithelial differentiation, we introduced a *SERPINB3*-containing lentiviral (or control) vector into primary basal cells from a single non-COPD donor at a multiplicity of infection (MOI) of 5-20 (**Table S5)**. We chose *SERPINB3* due to its high correlation with both *SERPINB4* and -*B13* in the HLCA and prior use in the literature (38). Hematoxylin and eosin (H&E) and AB-PAS staining of ALI-differentiated cells (D28) showed regions of apparent GCH as well as other areas containing cells with a flattened shape in the *SERPINB3* overexpression group (MOI=20) (**Fig. 4A and B**). To characterize these cells in more detail, we used SPLiT-seq(32) to perform scRNA-seq on ALI-differentiated cells (D28) across conditions (**Fig. 4C and D, Fig. S4C-E**). *SERPINB3* overexpression resulted in a condition-specific population of cells which expressed markers of goblet cells and inflammatory cytokines and chemokines including *CXCL1*, *CXCL2*, *CXCL3*, *CXCL5*, *CXCL6*, *CXCL8*, *CXCL16*, *CXCL17*, *CX3CL1*, and *CCL20* (**Fig. 4E, Fig. S4F, Table S6**). These cells were annotated as inflammatory goblet cells. Interestingly, we also observed an expansion of squamous basal and squamous cells in the *SERPINB3* overexpression group (MOI=20) (**Fig. 4E**).

**Fig. 4.**
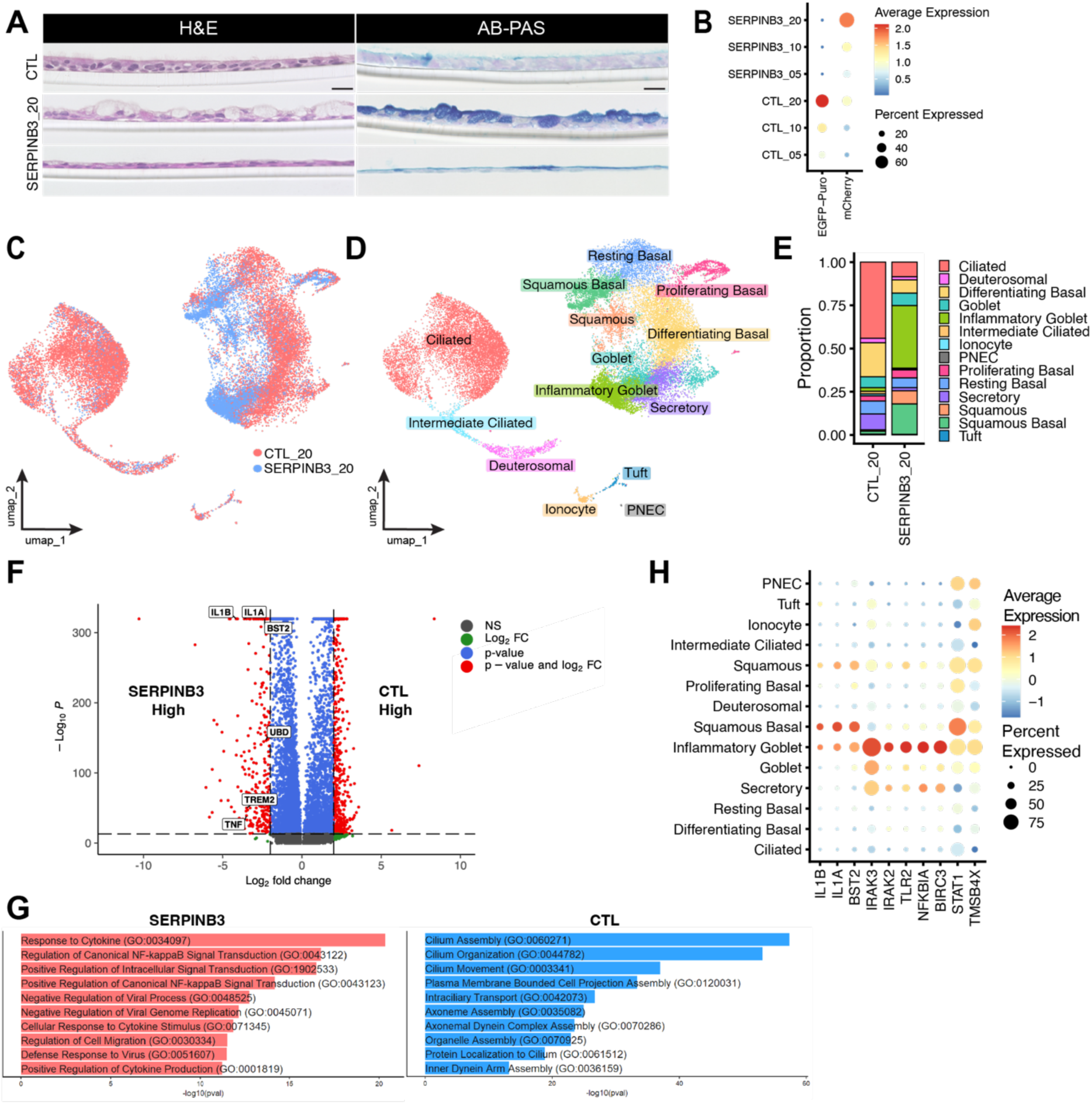
*SERPINB3* overexpression causes GCH and squamous cell metaplasia. **(A)** Representative images of ALI membranes in cross-section for each treatment group on ALI D28. Two regions were chosen for SERPINB3 overexpression at a multiplicity of infection (MOI) of 20 to display the varying phenotypes observed. Hematoxylin and Eosin (H&E) staining of cell morphology (left panel) and Alcian Blue-Periodic Acid-Schiff (AB-PAS) staining of mucin positive cells (right panel) Scale bar = 20 μM. **(B)** Dot plot showing absolute expression of the fluorescent reporters in control and SERPINB3 lentiviral constructs at each MOI dose. **(C)** UMAP of SERPINB3 and control overexpressing cells at 20 MOI colored by condition. **(D)** UMAP colored by annotated cell type in 20 MOI condition. **(E)** Proportion of cell types within each condition. **(F)** Volcano plot of differentially expressed genes in SERPINB3 20 MOI condition versus control 20 MOI condition across all cell types. *IL1B, IL1A, BST2, UBD, TREM2*, and *TNF* (highlighted) are significantly upregulated in the SERPINB3 20 MOI condition. **(G)** Enrichr comparison of GO Biological Process (2026) terms enriched in the SERPINB3 (left) or control (right) MOI 20 condition. **(H)** Dot plot showing select inflammatory gene expression across each cell type.

To evaluate the effects of *SERPINB3* overexpression on global gene expression, we compared gene expression across all cell types in the *SERPINB3* condition versus mock transfection using pseudobulking. Interestingly, *IL1A* and *IL1B* were among the top genes increased by *SERPINB3* overexpression (**Fig. 4F**). Enrichr analysis of DEGs showed enrichment in inflammatory terms including response to cytokine (GO:0034097) and regulation of canonical NF-KB signal transduction (GO:0043122) (**Fig. 4G**). Examination of genes within these GO terms by cell type indicated they were largely driven by inflammatory goblet cells, with a few increased in squamous basal cells (**Fig. 4H**). *IL1B* was expressed by inflammatory goblet as well as cells expressing squamous cell markers. Taken together, these data suggest that *SERPINB3* directly contributes to GCH and squamous cell metaplasia (SCM) and promotes expression of *IL1B* and other inflammatory cytokines and chemokines.

### A COPD-specific subset of basal cells express SERPINB4 and GC genes ex vivo

Since both squamous and goblet cells were increased by *SERPINB3* overexpression, we speculated that basal cell heterogeneity might account for these disparate differentiation outcomes. To investigate this further, we isolated, expanded, and performed RNA sequencing (RNA-seq) on 53 individual basal cell clones from cryopreserved tissue from five COPD lung explants and four non-COPD controls (**Fig. 5A, Table S7**). Hierarchical clustering of clones independent of sex and cell cycle influence indicated a COPD-specific cluster of seven clones with a distinct transcriptional profile that included expression of goblet cell genes including *MUC5AC*, *MUC5B*, *MSMB*, and *CEACAM5* (**Fig. 5B**). Although these clones were proliferating and expressed the basal cell markers *TP63*, *NGFR*, *KRT5*, and *ITGB4* (**Fig. 5C**), they were generally more transcriptionally similar to human lung goblet cells than basal cells (**Fig. 5D**).

**Fig. 5.**
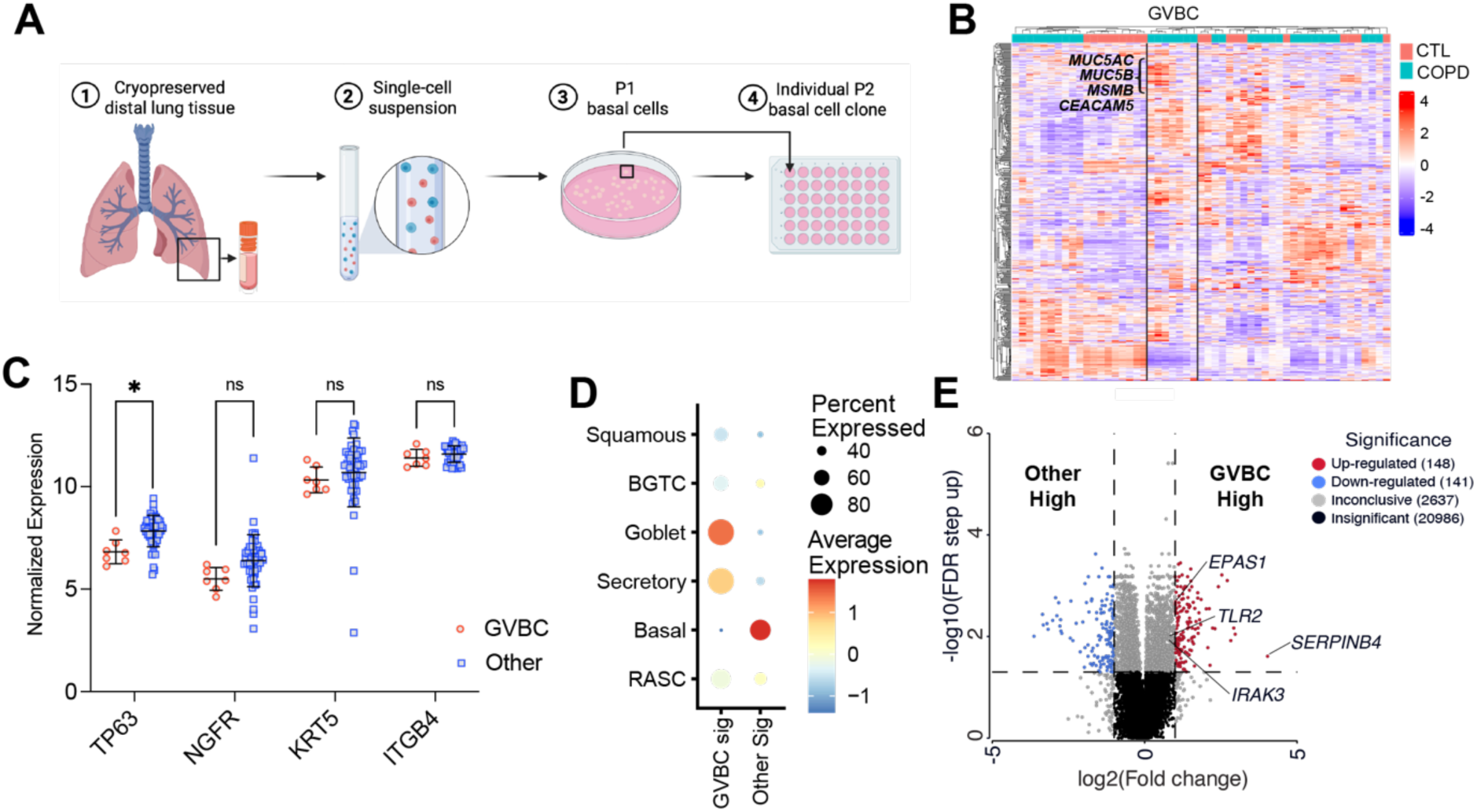
A *SERPINB4*+ population of COPD basal cells are transcriptionally similar to goblet cells. COPD and control basal cells were derived from distal lung tissue and expanded in culture to allow for individual clonal expansion. Discrete clones were lifted with cloning cylinders and expanded separately for downstream assays. **(A)** Graphic demonstrating the workflow for basal cell cloning. **(B)** Heatmap showing unsupervised hierarchical clustering on bulk RNA sequencing of 53 clones (19 control/ 34 COPD). A cluster of clones enriched for *MUC5AC, MUC5B, MSMB,* and *CEACAM5* were denoted as goblet-variant basal cells (GVBCs). **(C)** Expression of basal cell markers *TP63, NGFR, KRT5,* and *ITGB4* in GVBCs vs all other clones. *= FDR< 0.05; unpaired t-test **(D)** Dot plot of module scores based on the top 200 genes ranked by log_2_FC (and p<0.05) overlaid on the COPD SAEC object (**Fig. 1**). **(E)** Volcano plot showing differentially expressed genes in GVBCs versus all other basal cell clones. *SERPINB4* and *EPAS1* are among the significantly up-regulated genes (log_2_FC >2) in GVBCs. *TLR2* (log_2_FC =1.76) and *IRAK3* (log_2_FC =1.74) are also increased in GVBCs.

These findings were in contrast to all other clones from both COPD and control samples which were transcriptionally similar to basal cells from our human lung cell atlas (**Fig. 1**). We therefore called these cells “goblet-variant basal cells” or GVBCs. Notably, GVBCs were more transcriptionally similar to human lung goblet cells than BGTCs (**Fig. 5D**) indicating a more differentiated state. We compared gene expression of GVBCs to all other clones and found the top differentially expressed gene (DEGs) between these groups was *SERPINB4* (FC 16.3, FDR < 0.02) (**Fig. 5D**). Additionally, we found increased expression of *EPAS1* (encoding HIF-2A) (FC 2.02, FDR < 1.89 x 10^-3^), which has previously been shown to induce expression of clade B serpins(39), and *TLR2* (FC 1.76, FDR < 9.74 x 10^-3^) and *IRAK3* (FC 1.74, FDR < 0.01), which were among genes increased by *SERPINB3* overexpression in primary basal cells.

## DISCUSSION

This work advances the field by showing: 1) goblet cells primarily arise directly from basal cells in the COPD lung; 2) clade B serpins are transiently expressed during basal-to-goblet cell differentiation in the COPD lung and in vitro in response to cytokines; 3) *SERPINB3* overexpression in basal cells drives expression of inflammatory cytokines and expansion of inflammatory goblet, squamous and squamous basal cells; and 4) a COPD-specific population of GVBCs expresses high levels of *SERPINB4* as well as goblet cell markers. Together, these data provide a rationale for further study of clade B serpins as a therapeutic target to reverse GCH in COPD.

We describe a population of cells we term BGTCs with a transcriptional profile and physical location intermediate to basal and goblet cells, suggesting basal cells are the predominant precursor to goblet cells in COPD patients not experiencing an exacerbation. BGTCs are rare in control samples where they remain associated with goblet cells, but they are enriched in COPD lungs in regions of GCH. Similarly, we find that in response to IL-1β or IL-13 treatment goblet cells likely arise from basal cells through a distinct intermediate state. While the directionality of this transition state cannot be proven from gene expression data, the robust expression of metabolic genes in BGTCs is consistent with the high energy requirements for macromolecular synthesis typical of basal cell differentiation (40). These data do not preclude the possibility that other cell types can serve as goblet cell progenitors in the COPD lungs, as is known to occur in mice (41–43). Rapid transition states might not have been captured in our COPD scRNA-seq atlas, and none of the COPD patients whose samples were included in our scRNA-seq atlas were experiencing an exacerbation which could induce trans-differentiation of other epithelial cell types. Conversely, since control samples used in our scRNA-seq and spatial transcriptomic studies were from lungs rejected for organ donation, we cannot exclude the possibility that the few BGTCs observed in control samples were driven by pneumonia or other comorbid inflammatory conditions, incorrect annotation of smoking status, or confounding environmental exposures (44, 45)Additional studies will be required to quantify BGTCs in lung tissue from healthy non-smokers and in COPD patients experiencing an exacerbation.

We found that BGTCs were marked by high expression of multiple clade B serpins. In the HLCA, *SERPINB3* was expressed by club (nasal and non-nasal), goblet (nasal and bronchial), and hillock-like cells and along with *KRT5* expression was the canonical marker used to define suprabasal cells (46). We noted expression of *SERPINB3* in BGTCs which are physically located above basal cells within the airway epithelium, supporting the idea that BGTCs are a subset of suprabasal cells differentiating into goblet cells. BGTCs express a variety of genes related to cell adhesion and migration, further supporting the idea of their mobility within the airway epithelium. BGTCs express some markers of hillock-like cells from the HLCA including *RAB38* and *TNS4* (46), and *SERPINB2*, another clade B serpin, marks hillock luminal cells in mice (47). However, BGTCs lack the canonical hillock cell marker *KRT13*, and we did not observe BGTCs in hillock-like structures in our spatial transcriptomics atlas. Thus, shared expression of markers in BGTCs and hillock-like cells may reflect their shared status as transitional airway epithelial cell populations.

With the exception of *SERPINB1*, -*B6*, and -*B9*, all clade B serpins are located in a cluster on chromosome 18q21.3 (48). *SERPINB3*, -*B4*, and -*B13* are adjacent on chromosome 18q21.3, phylogenetically related, and share significant sequence homology (48). Using data from the HLCA and other atlases, we found expression of *SERPINB3*, *-B4*, and *-B13* to be highly correlated, suggesting a possible shared mechanism of regulation. *SERPINB3* and -*B4* have previously been shown to be induced by IL-4 and IL-13 in the airway epithelium (44), by TNF-α, IL-17A and IL-22 in keratinocytes (44, 49, 50), and by oncogenic RAS via MAPK signaling in IMR-90 fibroblasts (51). Additionally, *SERPINB3* has been shown to be upregulated by HIF-2α in a hepatocellular carcinoma cell line (39). In a study of large airway epithelial cells grown in ALI culture after isolation from human tracheas, multiple clade B serpins including *SERPINB3* and -*B13* were increased by cigarette smoke in various basal cell subpopulations (52). We found that IL-1β and IL-13 increased goblet cell formation and that cells differentiating into goblet cells expressed multiple clade B serpins including *SERPINB3*, *-B4*, and *-B13*. Increased expression was a shared feature of these transitioning cells despite transcriptional differences between those generated by IL-1β and IL-13. Although our study does not address the mechanism of regulation of clade B serpins, COPD is an inflammatory disease and increased expression of cytokines such as IL-1β could induce clade B serpin expression in nearby basal cells. We found upregulation of pathways related to aerobic respiration and the electron transport chain as a feature common to BGTCs and cells undergoing transition from basal to goblet cells in vitro following IL-1β treatment, supporting a potential role for IL-1β in generating this transitional cell population. Further studies will be required to understand how inflammation in the COPD lung microenvironment influences clade B serpin expression.

We found that *SERPINB3* overexpression in primary basal cells from a healthy donor increased numbers of squamous and squamous basal cells after differentiation in ALI. *SERPINB3* was originally called squamous cell carcinoma antigen 1 (SCCA1) after being isolated from human cervical squamous cell carcinoma (SCC)(53) and it has been used as a diagnostic biomarker for SCC of multiple organs (54). The emergence of squamous cells upon *SERPINB3* overexpression suggests it may contribute to SCC of the lung in addition to serving as a diagnostic biomarker. We also noted increased expression of *SERPINB2* in squamous cells from our scRNA-seq atlas, a gene previously used to identify murine hillock cells which are predisposed toward squamous cell differentiation (47) suggesting a possible mechanistic role in SCM. Finally, we noted increased numbers of squamous basal cells resulting from *SERPINB3* overexpression. While basal cell hyperplasia is another commonly observed feature of COPD (55, 56), it is unclear the extent to which squamous basal cells contribute to this phenotype.

Previous literature supports a mechanistic role for clade B serpins in goblet cell formation. Genetic deletion of *Serpin3a*, a mouse homolog of human *SERPINB3/4*, abrogates house dust-mite and IL-13-induced GCH in mice (57) and serum levels of SERPINB3/4 (assayed together) have been shown to be increased in the serum of patients with asthma (44). We found *SERPINB3* overexpression leads to expression of a variety of inflammatory genes and cytokines, including a small population of inflammatory goblet cells expressing IL-1β. In this manner, *SERPINB3* may serve as a local amplifier of inflammatory cytokine signaling within the airway epithelium. Clade B serpins are “suicide-substrate”-like protease inhibitors that are thought to predominantly function intracellularly (48), though they are detected in serum in multiple disease contexts. The specific mechanism through which this class of clade B serpins activates inflammatory signaling is an important area for future study. Although *SERPINB3* and -*B4* share significant sequence homology, -*B3* targets cathepsins K, L, S, and V while -*B4* targets cathepsin G and chymase (48). These specificities could offer important clues as to the mechanism of action of clade B serpins in basal cell differentiation. Regardless of the mechanism of action, *SERPINB3* and/or *-B4* may serve as novel biomarkers of GCH in COPD as has been observed in asthma and various malignancies. Future studies that correlate serum levels of *SERPINB3/4* with radiographic biomarkers of GCH (i.e. mucus plugging) would be helpful.

We observed a small number of COPD-specific basal cell clones which were more transcriptionally similar to goblet cells than basal cells. We termed these cells “goblet-variant basal cells” or GVBCs. These findings are consistent with other publications indicating some COPD basal cells are primed toward goblet cell differentiation (22, 27, 28). GVBCs displayed increased expression of *SERPINB4* and *EPAS1*. Prior studies using hepatocellular carcinoma cell lines indicate a mutually reinforcing relationship between hypoxia signaling and clade B serpin expression (39, 58). Additional research will be required to determine the relationship between hypoxia and clade B serpin expression in COPD airways and the epigenetic mechanism(s) responsible for persistently increased expression of *SERPINB4* in COPD basal cells.

In summary, this work implicates basal cells generally, and clade B serpins specifically, as causal features of GCH in the COPD lung. Therefore, clade B serpins could be an important therapeutic target to reduce GCH in COPD.

## MATERIALS AND METHODS

### Human lung tissue acquisition and approvals

Explanted lungs from COPD patients undergoing lung transplantation were obtained after informed consent according to an institutional review board (IRB)-approved protocol from Vanderbilt University Medical Center (IRB #060165). Control samples were obtained from deceased organ donors and exempted from IRB review. Demographic and clinical information for samples undergoing scRNA-seq has been previously described (30).

### Human single-cell lung atlas

The COPD epithelial data set was composed and processed as previously described (30).

Briefly, raw counts were processed by Cell Ranger (v6.1.2). Poor quality cells were removed, then data was normalized, and variable genes were determined utilizing RISC (Robust Integration of Single Cell RNA-seq data) (v1.6.0). Integration of GSE196341, GSE168191, and GSE178360 with additional samples enriched for epithelial cells was performed using Reference Principal Component Integration. Seurat v4 was used for post-integration processing and analysis. Following annotation utilizing the LungMap web-based Azimuth tool and assisted by canonical markers, airway epithelial cells were isolated for further analysis.

Basal and goblet cells from the small airway epithelial object described above were subclustered using Seurat v5 and 12 principal components (PCs, chosen after examining an elbow plot). A clustering resolution of 0.2 was chosen based on ability to resolve multiple clusters while avoiding donor-specific grouping. This resolution yielded three goblet clusters, two basal cell clusters, a cluster of rare cells, and a cluster of cells that were originally annotated as goblet cells but also expressed the basal cell markers *KRT5*, *TP63*, and *ITGB4* (BGTCs). The top 50 markers were determined by running FindMarkers with the percent cut off set to 0.25 then running top n for the BGTC cluster markers using average log2 fold change.

Gene set enrichment analysis (GSEA) was performed on BGTC positive and negative markers calculated using FindMarkers with the percent cut off set to 0.25 and a log2 fold change threshold of 0.5. GSEA for gene ontology was performed using the R package clusterProfiler(59) with the following settings: ont =“BP”, keyType = “SYMBOL”, minGSSize = 10, maxGSSize = 800, pvalueCutoff = 0.05, OrgDb = ’org.Hs.eg.db’, nPermSimple = 100000, eps = 0, scoreType = “std”.

### Tissue array construction and spatial transcriptomics

Tissue arrays were constructed in a 3 x 1 configuration of 5 x 5 mm squares of tissue from non-diseased deceased lung donors (controls) and COPD explants. Sections were taken from formalin fixed, paraffin-embedded (FFPE) samples using 5 x 5 mm square punches and re-embedded into a row of 3 with one region of interest (ROI) isolated from a control sample and the remaining two ROIs from two separate lobes of a COPD explant. 5 μm thick tissue sections were placed onto Xenium slides and sent to the 10x Genomics Catalyst team for processing. The human lung panel and a custom panel containing 50 BGTC markers along with 50 other genes related to stem cell function and goblet cell formation was utilized (Design ID: DNKKBE).

Multitarget IF was utilized for robust cell segmentation and an H&E was performed and imaged following the Xenium assay to compare morphology of the cells to expression of canonical markers. Samples were analyzed utilizing Seurat v5 and the Xenium Explorer 3 application for visualizations. SCTransform was utilized to normalize data. 30 PCs were chosen based on elbow plot and genes, and a resolution of 0.4 was chosen for Leiden clustering. Annotations were performed based on expression of marker genes. Airway epithelial cells were isolated and reclustered for finer annotation. 20 PCs were utilized and Leiden clustering with a resolution of 0.6 was performed. This yielded seven final clusters. Markers were calculated utilizing Seurat’s FindMarkers function with only positive markers and a logFC threshold of 0.1. Top markers using inverse adjusted p-value were found for each cluster to confirm accuracy of annotations.

### Human small airway epithelial cell (HSAEC) culture and cytokine exposure

#### Culture

Human Small Airway Epithelial Cells (HSAECs; ATCC, #PCS-301-010) were initially expanded (P2-P3) in 10 cm dishes in PneumaCult-Ex Plus Medium (Stemcell Technologies, #05040) with Penicillin Streptomycin (P/S, 1:100, Corning, #30-002-Cl) and then 80,000 cells were seeded in Transwell membranes (6.5mm, Costar, #3470). At 70% confluency they were transitioned to Air Liquid Interface (ALI) using PneumaCult-ALI for Small Airway media (Stemcell Technologies, #05050), marking ALI day 0 (D0). The apical side of the membrane was washed every other media change with PBS for 5 minutes at 37℃.

#### Cytokine exposure

On ALI days 14-27, cytokine treatment was added to the basolateral chamber during each media change [IL-13 10 ng/mL (PeproTech, #200-13) or IL-1B 10 ng/mL (PeproTech, #200-01B) or vehicle control (1:1000 0.1% BSA)].

#### SPLiT-pool ligation-based sequencing

On ALI D29, cells were lifted from membranes with the ACF Cell Dissociation Kit (Stemcell Technologies, #05426). They were fixed according to the Evercode Cell Fixation v2 User Manual v2.0.1 and prepared for sequencing using the Evercode WT v2 User Manual (Parse Biosciences; Seattle, WA). Next-generation sequencing was performed on an Illumina NovaSeq 6000 (S4) PE150 at Vanderbilt Technologies for Advanced Genomics (VANTAGE). The Parse Biosciences Software (Pipeline V1) was used to de-multiplex and align (Human GRCh38) FASTQ files, then generate a filtered digital gene expression (DGE) matrix using default parameters. The DGE matrix was the input to create a Seurat Object (v4.3) in R (v4.3.0). Quality control criteria excluded outliers with >10,000 genes, >40,000 transcript counts, and >20% of genes mapping to mitochondrial UMIs. The standard Seurat workflow was run (R v4.5.2, Seurat v5.3.0): NormalizeData (LogNormalize), ScaleData (all genes), RunPCA, FindNeighbors (dims = 1:20), FindClusters (Leiden, resolution= 0.4), RunUMAP (dims = 1:20). Cluster markers were identified using the FindAllMarkers command (min.pct = 0.25 and logfc.threshold = 0.25). Cell type annotation was based on canonical markers. Basal and secretory cells were isolated (33,936 cells) and reclustered as follows (R v4.5.2, Seurat v5.3.0): FindNeighbors (dims = 1:20); FindClusters (Leiden, resolution= 0.5); RunUMAP (dims = 1:15). Markers were identified using the FindAllMarkers command (min.pct = 0.25 and logfc.threshold = 0.25). The top 5 markers for each cluster were identified after filtering for p < 0.05 and ranking by logFC.

#### Monocle3

Monocle3 (33, 34) was utilized with R version 4.5 and Bioconductor version 3.14. SeuratWrappers was used to run Monocle3 directly on the Seurat objects. Partitions were identified then the partition containing basal and goblet/secretory cells was isolated and trajectory analysis was run. Resting basal cells were selected as the root node. Trajectory analysis predicted three separate trajectories (IL-1B induced, IL13-induced, and control secretory). Each of these branches were selected and assessed individually for gene expression of SERPINB3, 4, and 13 over pseudotime.

### HSAEC culture and SERPINB3 lentivirus overexpression

#### Culture and Lentiviral Transduction

HSAECs (ATCC, PCS-301-010) were initially grown in PneumaCult-Ex Plus Medium (Stemcell Technologies, #05040) with P/S. Cells were lifted and treated in suspension for 3 hours at 37℃ at 5, 10, or 20 MOI with CTL (VectorBuilder, #VB010000-9492agg) or *SERPINB3* (VectorBuilder, #VB251006-1466ftu) containing lentivirus and polybrene (5 µg/mL). 80,000 cells were seeded on Transwell membranes (6.5mm) in Ex-Plus media. At greater than 70% confluency they were transitioned to ALI culture and PneumaCult-ALI-S media (Stemcell Technologies, #05050) replaced the media in the basolateral chamber. The apical chamber was washed every other media change with PBS for 5 minutes at 37℃.

#### SPLiT-pool ligation-based sequencing

On ALI D28, the cells were dissociated using the ACF Cell Dissociation Kit (Stemcell Technologies, #05426), then fixed with Evercode Cell Fixation v4 (Parse Biosciences). Fixed cells were prepared for sequencing using the Evercode WT kit v4 (Parse Biosciences). Next-generation sequencing was performed on an Illumina NovaSeq 6000 (S4) PE150 at VANTAGE. Trailmaker (Pipeline v1.7.3, Parse Biosciences) was used to de-multiplex and align (Human GRCh38 and custom genes for mCherry and EGFP-puromycin) FASTQ files. The filtered DGE matrix from the pipeline output was used to create a Seurat Object (v5.5) in R (v4.4.3). Quality control criteria excluded outliers with >5,000 genes, >10,000 transcript counts, and >20% of genes mapping to mitochondrial UMIs. Seurat was used to normalize (Log Normalize) and scale the object. The object was clustered by the following parameters: RunPCA; FindNeighbors (dims = 1:30); FindClusters (Louvain, resolution= 0.5); RunUMAP (dims = 1:30). Cell-type annotation was based on canonical marker expression and using the FindAllMarkers command (min.pct = 0.25 and logfc.threshold = 0.25). Cells containing high levels of counts and features were excluded as doublets and the above parameters were repeated. The top 5 markers for each cluster were identified after filtering for p < 0.05 and ranking by logFC. A 20 MOI object was created for targeted analysis.

### Human lung basal cell isolation and culture

#### Cryopreservation

Lung tissue was collected in Dulbecco’s Modified Eagle Medium (DMEM, Gibco, #A1443001) and kept at 4℃ until being minced with scissors. Tissue was cryopreserved in 1 g aliquots with 500 μL of freezing media made of 90% FBS and 10% DMSO.

#### Culture

Cryopreserved tissue was briefly thawed and transferred to 20 mL of ice cold DMEM to dilute freezing media. One gram of minced tissue was transferred to 5 mL of enzyme solution (Multi Tissue Dissociation Kit 1, Miltenyi Biotec, #130-110-201) in a gentleMACS C Tube (Miltenyi Biotec, #130-093-237). The tissue was dissociated using a gentleMACS Octo Dissociator and then transferred to ice. 10 mL of DMEM was added to each C Tube and then cell suspensions were serially filtered through gauze then 100 μM and 40 μM cell strainers. Cells were resuspended with DNase 1 (Millipore, 5MU/mL) for 5 minutes at room temperature and then in ACK Lysing buffer for 3-5 minutes. Single-cell suspensions were seeded onto 10 cm dishes for expansion of basal cells in PneumaCult-Ex Plus Medium (Stemcell Technologies, #05040) with Cipro (10 μg/ml), Fungizone (1:100, Gibco, #15290018), and P/S (1:100) for 2-3 days. Then Ex Plus media with P/S was replaced every other day for the duration of culture.

#### Clone isolation

To allow for adequate space for individual basal cell clones to expand, 200,000 live cells from the single-cell suspensions described above were seeded on 10 cm dishes.

Individual basal cell clones expanded for 12-15 days in culture to become a sufficient size for cloning without merging with other clones. Before cloning, the media was removed and the dish was washed twice with PBS. 10 x 10 mm cloning cylinders (PYREX #3166-10) were plunged into sterile grease (DOW Corning High Vacuum Grease, Thermo Scientific, #044224.KT) and then secured around individual colonies of basal cell clones. ACF dissociation solution (Stemcell Technologies, #05427) was added to individual cylinder wells and incubated at 37℃ for 15 minutes. Then, an equal part of ACF inhibition solution (Stemcell Technologies, #05428) was added to each well. Isolated clones were transferred to unique tubes and resuspended in Ex Plus media to be seeded at passage 2 for further expansion on 12-well plates coated with collagen (Human collagen from placenta, 6 μg/cm; Sigma Aldrich).

#### RNA-sequencing

At 80% confluence, P2 basal cell clones were lifted (ACF Cell Dissociation Kit, Stemcell Technologies, #05426), washed in PBS, and collected for RNA using a RNeasy Mini Kit (Qiagen, #74104) according to manufacturer recommendations. Samples were sequenced (Novogene Corporation Inc.) on an Illumina NovaSeq 6000 (Paired-End (PE) 150 bp) targeting 20 million raw read pairs per sample. Partek Flow (Illumina) was used to align reads with HISAT2, normalize (CPM, Add 1, Log 2), and adjust counts for batch effect (General linear model). For the creation of the heatmap, unsupervised hierarchical clustering was performed in iDEP(60) (v2.01) using normalized (Rlog) and batch effect adjust counts (General linear model). ***Module scores***. Seurat’s module score function was used to draw comparisons between datasets using the top 200 significantly enriched genes (sorted by log-fold change) for the cell type being compared.

### RNA In Situ Hybridization and Immunofluorescence

#### Tissue

##### RNA-In Situ Hybridization

RNA-ISH was performed utilizing the RNAscope Multiplex Fluorescent Reagent Kit v2 from ACD Bio/Biotechne according to the manufacturer’s instructions. A probe targeting *SERPINB4* (1050531-C3), which also detects *SERPINB3* due to high homology, was utilized. Following the RNAscope protocol, samples were washed 2x with TBST then blocked with TBS + 1% BSA for 30 minutes. Primary antibodies targeting KRT5 (Biolegend, #905903, 1:500) and MUC5B (Sigma Aldrich, #HPA008246, 1:100) were then applied to the tissue overnight at 4℃. Samples were then washed 2x and incubated with the appropriate secondaries (DAC-488 (1:100, Jackson ImmunoResearch Labs, # 703-545-155; DAR-cy3 (1:100, Jackson ImmunoResearch Labs, #711-165-152) for 45 minutes followed by 2 washes then incubation with DAPI (1:1000, Invitrogen, #D1306) for 10 minutes for nuclear visualization. Samples were then washed again and mounted with Prolong Gold mounting media and cured overnight before imaging. Samples were imaged using a Keyence BZ-X710 all-in-one fluorescence microscope with the appropriate filter cubes.

##### KRT5 and MUC5B

Samples were deparaffinized then washed twice prior to antigen retrieval in citrate buffer (pH6) for 20 minutes. Samples were then cooled, washed twice and treated with 0.1% triton for 5 minutes, then washed twice and blocked with 1% BSA in PBS for 30 minutes followed by staining as written above.

#### Cell culture

On ALI D28 of the cytokine and overexpression model, Transwell membranes were washed twice with PBS and then fixed in 4% paraformaldehyde (PFA) for 20 minutes. The membranes were removed from the Transwell inserts and cut into strips for embedding and sectioning. All images were processed in Fiji Image J using the same settings within stains. *Hematoxylin and Eosin (H&E).* Mayer’s Hematoxylin (Electron Microscopy Sciences, #26043-06) and Eosin-Phloxine Stain Set (Newcomer Supply, #1082A) were used for standard H&E staining. 40x images were taken using a Keyence BZ-X800 microscope.

##### Alcian Blue-PAS

Slides were deparaffinized and then stained according to manufacturer’s instructions of the Alcian Blue/PAS stain kit (Newcomer Supply, #91022B). Images were taken at 40x using a Keyence BZ-X800 microscope.

##### MUC5AC

Slides were deparaffinized and then incubated in 0.2% Triton for 5 minutes. The slides were blocked with 1% BSA for 10 minutes and then incubated with MUC5AC [1:100, Sigma Aldrich, #MAB2011 (clone CLH2)] overnight at 4℃. The next day the slides were incubated with Alexa Fluor 488 (1:100, Jackson ImmunoResearch Labs, #715-545-150) for 45 minutes at room temperature. DAPI (1:1000, Invitrogen, # D1306) was added for 10 minutes before cover slipping with Prolong Gold AntiFade Reagent (Invitrogen, #P36930). Images were captured using a Keyence BZ-X800 fluorescence microscope with the 40x objective.

##### MUC5B

Slides were deparaffinized and then incubated in 0.2% Triton for 5 minutes. The slides were blocked with 1% fish gelatin (Sigma Aldrich, #G7041) for 10 minutes and then incubated with MUC5B (1:100, Sigma Aldrich, #HPA008246) for 3 hours at room temperature. Then slides were incubated with Alexa Fluor Cy3 (1:100, Jackson ImmunoResearch Labs, #711-165-152) for 45 minutes at room temperature. DAPI (1:1000) was added for 10 minutes before cover slipping. Images were captured by confocal (Zeiss LSM 980) using the 63x objective and 3×3 tiling.

### Artificial Intelligence

#### Human lung cell atlas analysis in LungChat

##### Dataset and Preprocessing

Analyses were performed in LungChat (61) using the Human Lung Cell Atlas (HLCA; Sikkema et al., *Nature Medicine*, 2023), accessed via its integrated single-cell transcriptomic reference comprising 2,282,447 cells from 486 individuals across 49 datasets.

Following the HLCA processing pipeline, cells were aggregated into “supercell” metacell bins of 50 cells each (computed at the donor and cell-type level), yielding 50,520 metacells used as the unit of analysis. The HLCA core (107 individuals, healthy lung) was integrated using scANVI, and the extended HLCA (35 additional datasets spanning multiple lung diseases) was mapped onto the core reference using scArches, with cell-type annotations transferred accordingly. Gene expression values used in downstream analyses were normalized, log-transformed metacell-level counts (raw counts are also available in the source object). Cells annotated as “Basal resting” in the HLCA cell-type ontology were selected as the population of interest.

##### Gene Co-expression Analysis

For each query gene (*SERPINB3*, *SERPINB4*, *SERPINB13*), Pearson correlation coefficients were computed between the query gene and all other detected genes across metacells within the Basal resting cell population. Pairwise correlations among the three query genes themselves were also computed directly (*SERPINB3*–*SERPINB4*: r = 0.778; *SERPINB4*–*SERPINB13*: r = 0.696; *SERPINB3*–*SERPINB13*: r = 0.627). Genes with Pearson r > 0.4 relative to each query gene were retained as a co-expression gene set (*SERPINB3*: 305 genes; *SERPINB4*: 379 genes; *SERPINB13*: ∼90 genes), reflecting a moderate-to-strong positive co-expression threshold.

##### Overlap Analysis

Pairwise and three-way overlaps among the *SERPINB3*, *SERPINB4*, and *SERPINB13* co-expression gene sets (r > 0.4) were determined by set intersection and visualized as a Venn diagram, identifying a shared 67-gene “core module” common to all three gene sets.

## Acknowledgements

The authors wish to thank Dr. Vasiliy Polosukhin and Dr. Susan Guttentag for a critical review of the manuscript. Additionally, the authors would like to acknowledge the Vanderbilt Technologies for Advanced Genomics core (VANTAGE) at Vanderbilt for support with single-cell sequencing, split-pool sequencing, and RNA-sequencing; and the 10X Xenium Catalyst team for raw data acquisition and slide staining for the COPD Xenium data.

## Funding

American Thoracic Society/GlaxoSmithKline Research Grant in Obstructive Lung Disease (J.B.B)

National Institutes of Health grant 5T32HL094296 (J.B.B./ T.S.B.)

Department of Veterans Affairs grant IK2BX003841 and I50CX002551 (B.W.R.) A charitable gift from the Ann Duffer Family Foundation (B.W.R.)

## Author Contributions

Conceptualization: T.S.T, J.B.B, B.W.R

Methodology: T.S.T, J.B.B, D.S.N, C.M.S, B.W.R

Investigation: T.S.T, J.B.B., D.S.N., A.V., B.W.R

Visualization: J.B.B, T.S.T, B.W.R

Supervision: T.S.B, B.W.R

Writing—original draft: J.B.B, T.S.T, B.W.R

Writing—review & editing: T.S.T, J.B.B, C.M.S., L.B.W, B.W.R

## Competing Interests

The authors declare that they have no competing interests.

## Data, code, and materials availability

Data needed to evaluate the conclusions in the paper are present in the paper and/or the Supplementary Materials. Upon publication, raw sequencing reads and limited metadata will be deposited on the NCBI Gene Expression Omnibus (GEO) repository.

## Notes

### Competing Interest Statement

The authors have declared no competing interest.

