## Supplemental Figures for "Clade B serpins promote direct basal-to-goblet cell differentiation in airways of patients with chronic obstructive pulmonary disease"

### 1 Supplementary Figures

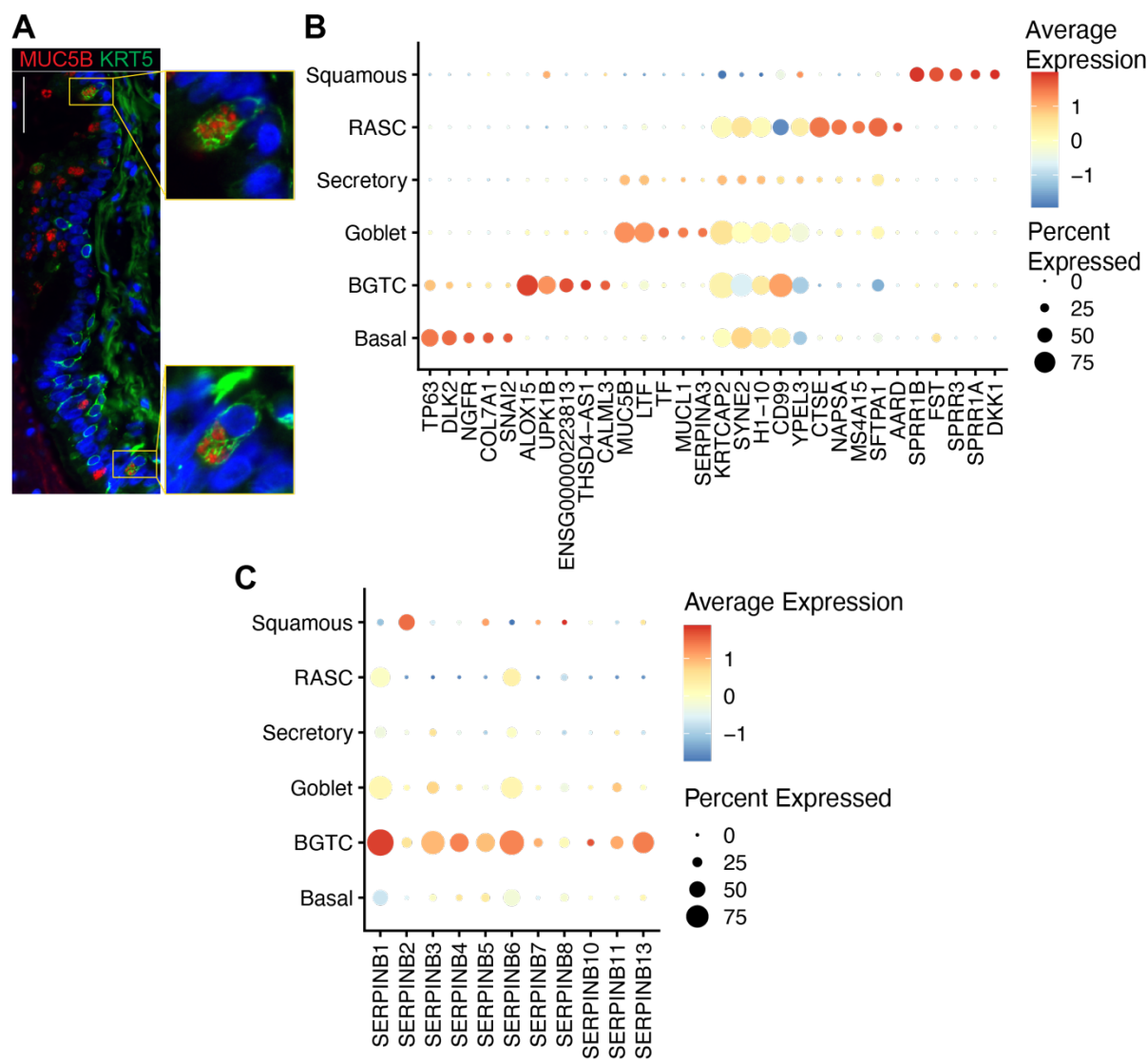

**Fig. S1.** (A) Immunostaining of KRT5 (green) and MUC5B (red) in a COPD airway with MUC5B+ goblet cells. Callout is 3x magnification. Scale bar = 50µm. (B) Top 5 markers of SAECs ranked by p-value (adjusted $p < 0.05$) then logFC (log FC > 0.25). (C) Dot plot of clade B Serpin expression in SAECs.

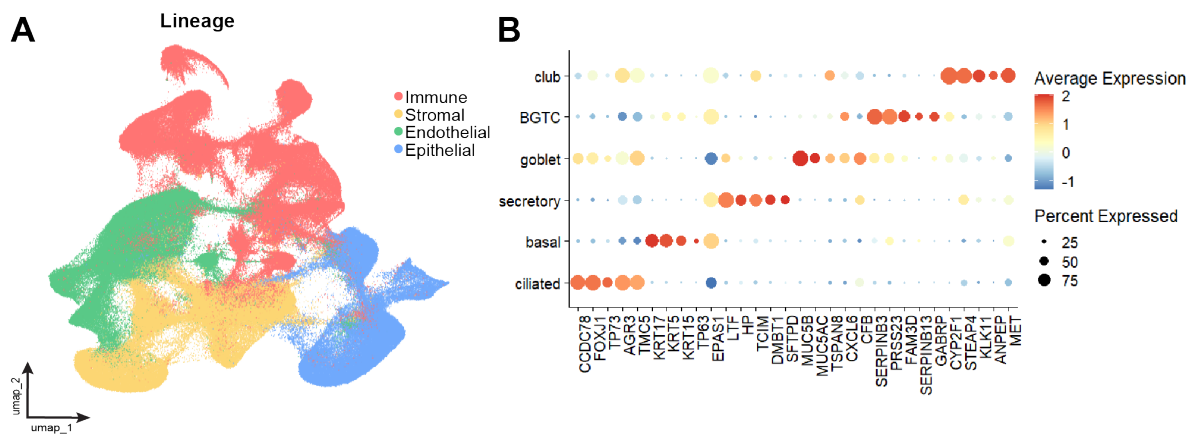

**Fig. S2. (A)** UMAP of all cells obtained using the Xenium Lung panel and BGTC markers with lineages labeled. **(B)** Dot plot of the top 5 makers of each cell type for the COPD and control Xenium airway object.

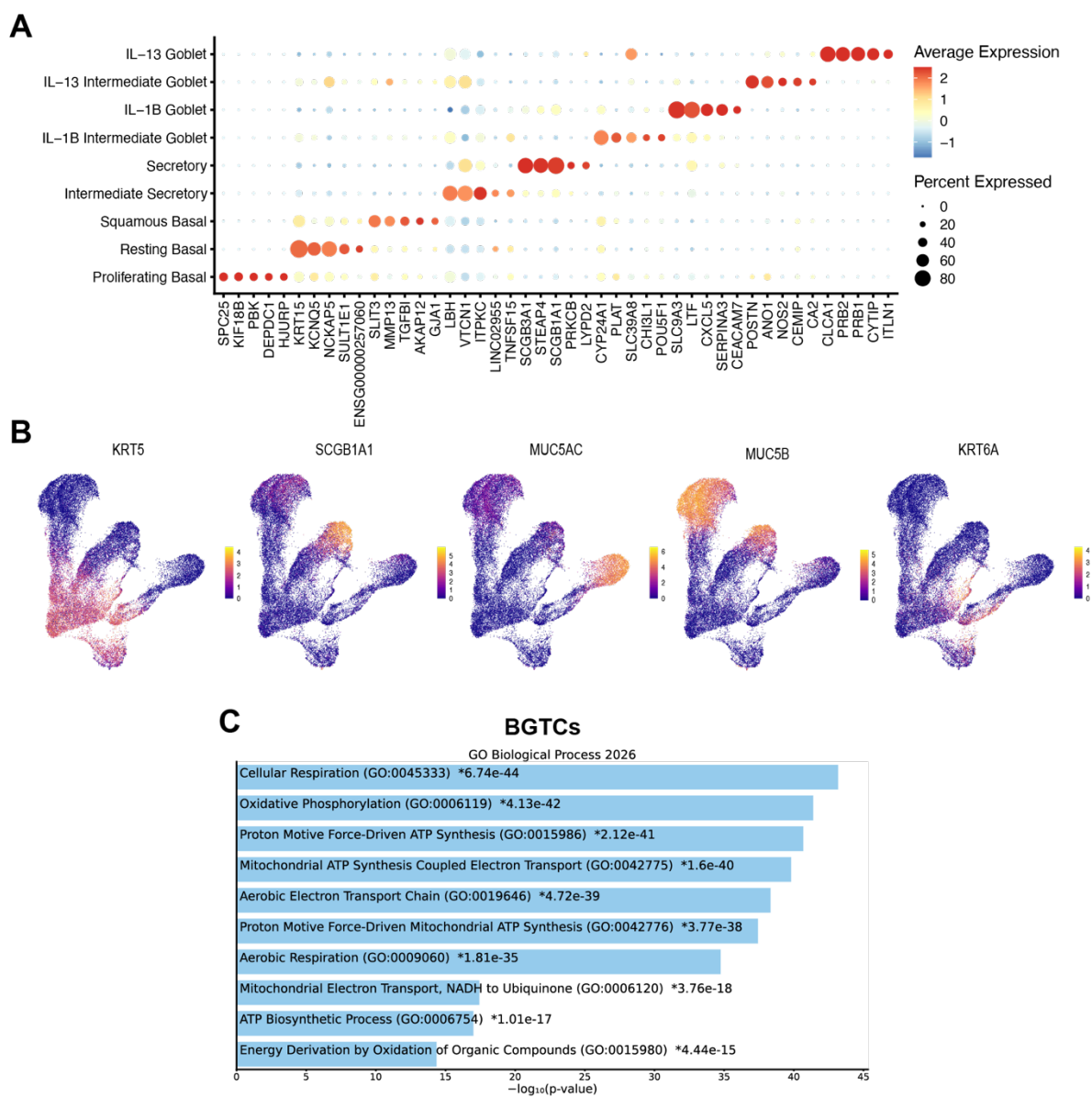

**Fig. S3. (A)** Dot plot of the top 5 markers for each cell type ranked by $\log FC > 0.25$ and $p < 0.05$. **(B)** Feature plots displaying showing expression of *KRT5*, *SCGB1A1*, *MUC5AC*, *MUC5B*, and *KRT6A* on the UMAP. **(C)** Enrichr bar plot based on cell type markers for BGTCs (**Fig. 1**) showing top GO Biological Process (2026) terms.

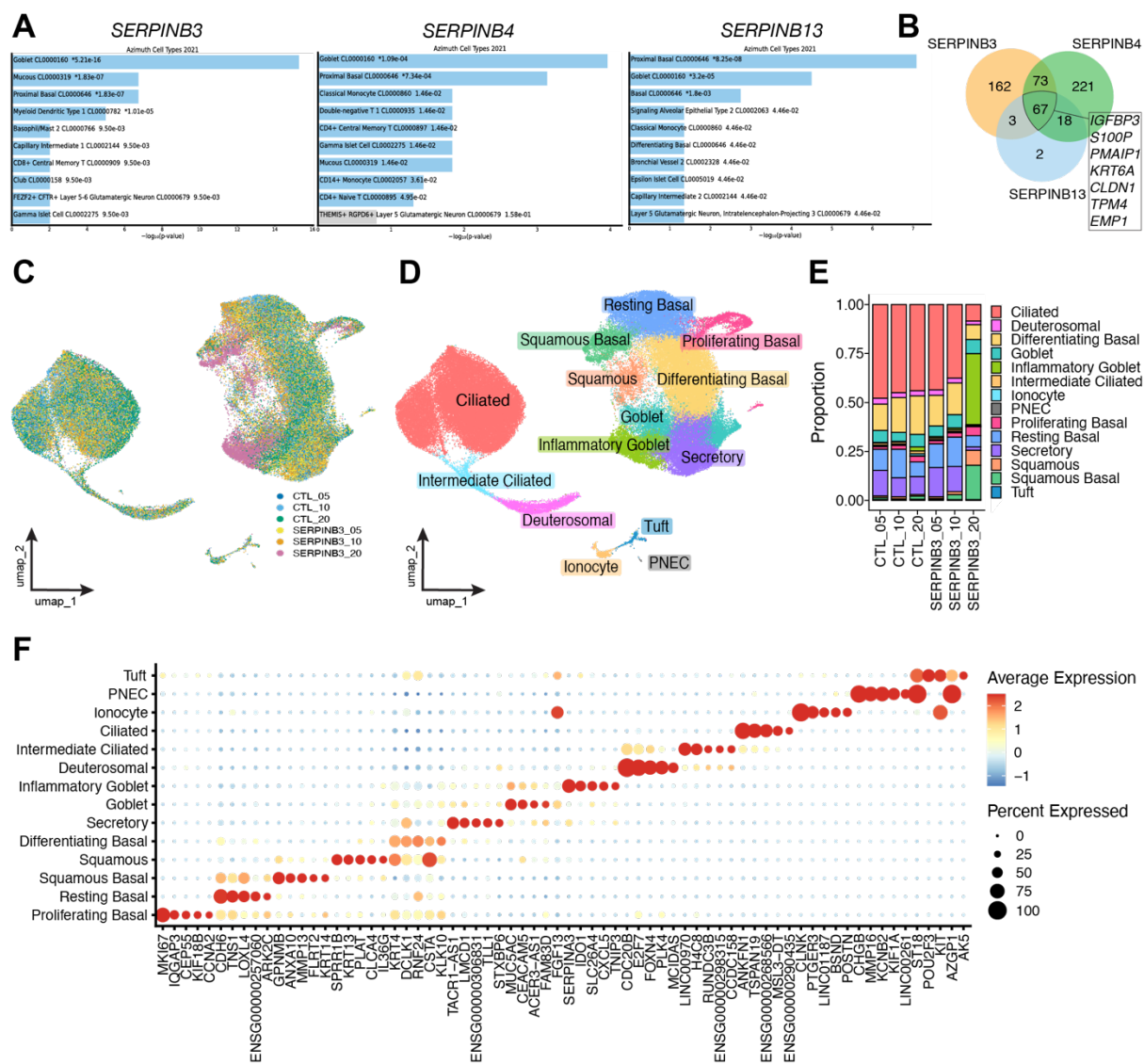

**Fig. S4.** (A) Enrichr bar plots displaying the top ten matches using the Azimuth Cell Types (2021) for genes correlated with *SERPINB3*, *SERPINB4*, or *SERPINB13* in the Human Lung Cell Atlas dataset (Pearson $r > 0.4$) (B) Venn diagram showing overlap of genes correlated with *SERPINB3*, *SERPINB4*, and *SERPINB13*. (C) UMAP of *SERPINB3* and control overexpressing cells colored by condition. (D) UMAP colored by annotated cell type for all conditions. (E) Proportion of cell types within each condition. (F) Dot plot of top 5 markers for each cell type.
